## Supplementary Figures 1-5 for "Multiplets in scRNA-seq data: extent of the problem and efficacy of methods for removal"

A Preprint

Dimitris Ttoouli*[1] and Daniel Hoffmann[1]
[1]Bioinformatics and Computational Biophysics, University of Duisburg-Essen, Essen, Germany

June 9, 2025

### Supplementary figures

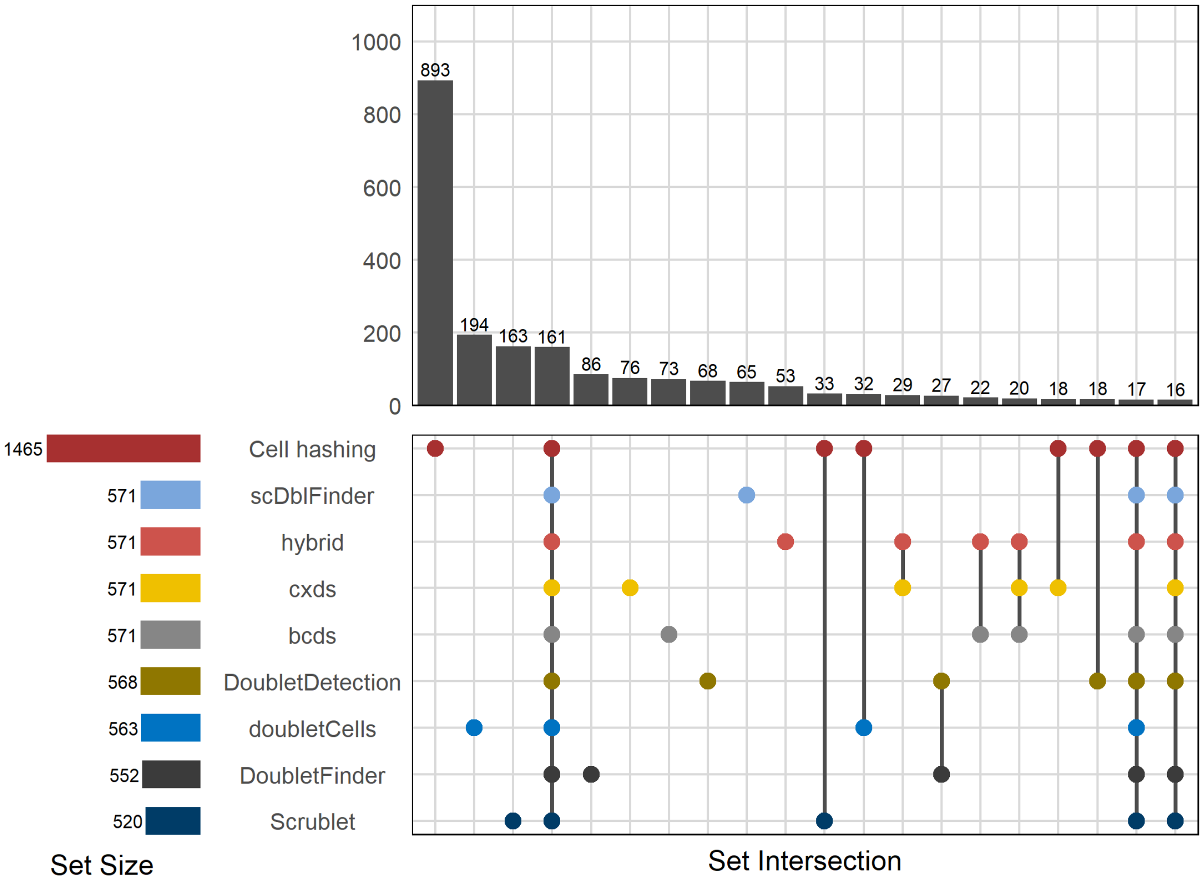


**S1 Fig.**
**Top 20 multiplet sets and intersections (cline dataset)**
UpSet plot showing sets and set intersections of multiplets identified by the tools and by cell hashing in the cline dataset. Horizontal bars (bottom left) show the total number of multiplets identified by each method. Vertical bars (top right) show the sizes of the sets of detected multiplets. If a point is present for a method underneath a bar (bottom right

*

plot panel), this means that this method has identified the particular multiplets that are members of the set denoted by that bar. If multiplet points are present underneath a bar, this means that multiple methods identified the same set of multiplets, i.e. their sets of multiplets intersect. Points are colored by method. Only the 20 largest multiplet sets and set intersections are shown. Note the y axis break.

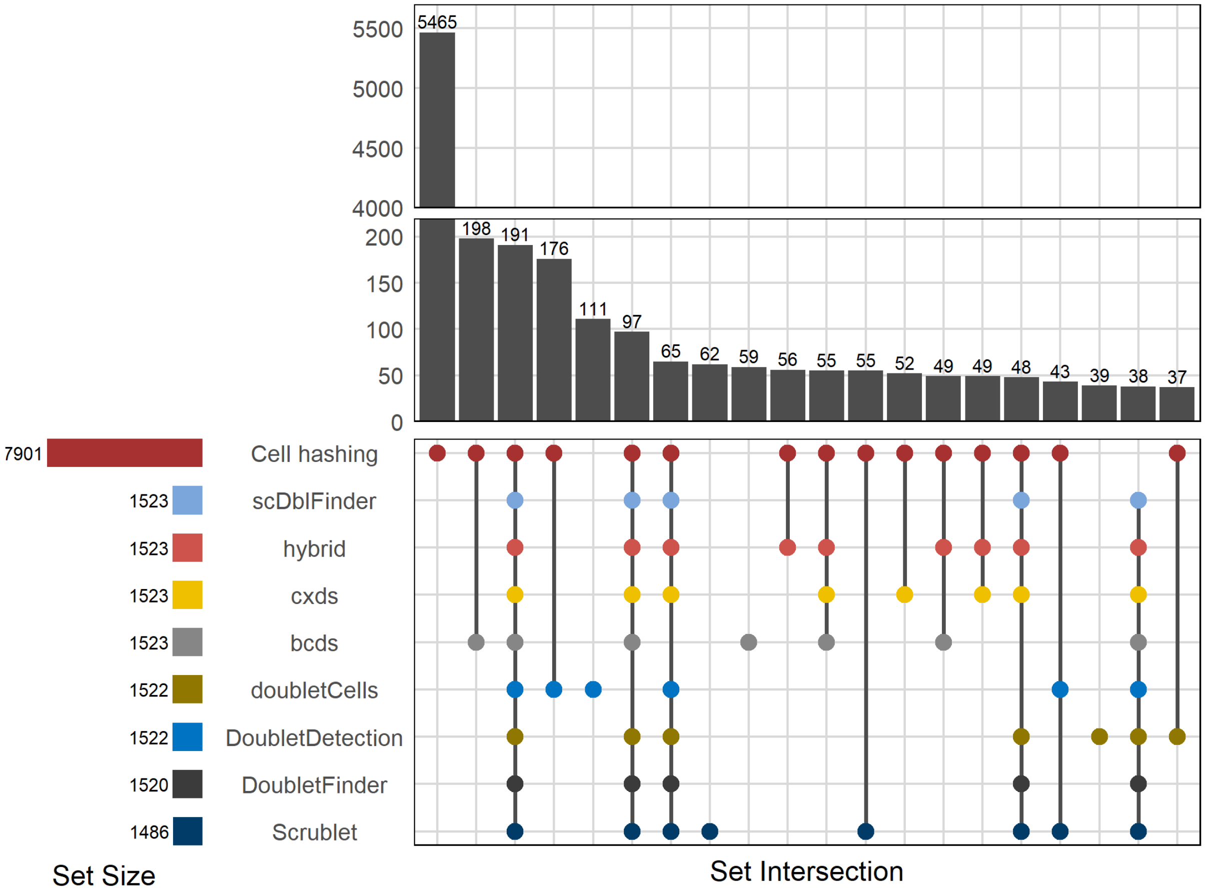


**S2 Fig.**

**Top 20 multiplet sets and intersections (mkidney dataset)**

UpSet plot showing sets and set intersections of multiplets identified by the tools and by cell hashing in the mkidney dataset. Horizontal bars (bottom left) show the total number of multiplets identified by each method. Vertical bars (top right) show the sizes of the sets of detected multiplets. If a point is present for a method underneath a bar (bottom right plot panel), this means that this method has identified the particular multiplets that are members of the set denoted by that bar. If multiplet points are present underneath a bar, this means that multiple methods identified the same set of multiplets, i.e. their sets of multiplets intersect. Points are colored by method. Only the 20 largest multiplet sets and set intersections are shown. Note the y axis break.

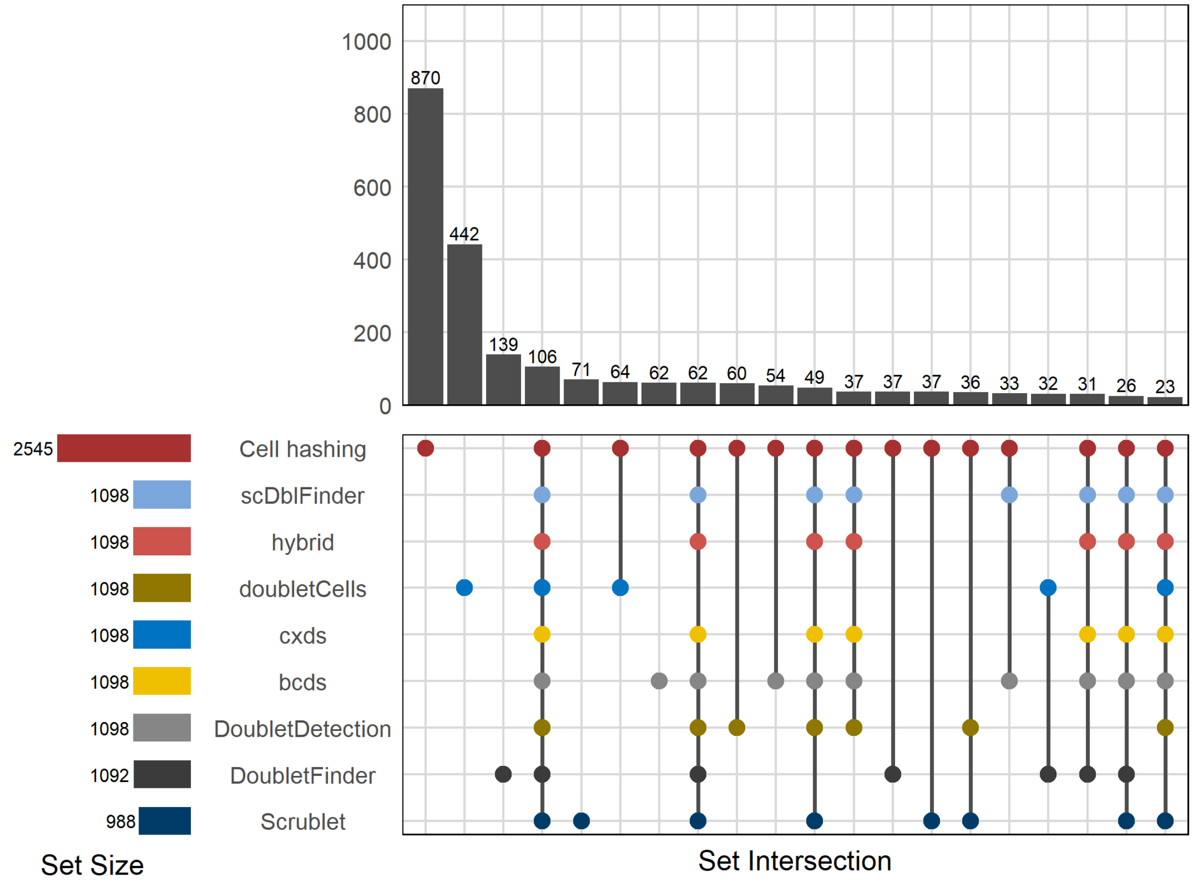


**S3 Fig.**

**Top 20 multiplet sets and intersections (pbmc dataset)**

UpSet plot showing sets and set intersections of multiplets identified by the tools and by cell hashing in the pbmc dataset. Horizontal bars (bottom left) show the total number of multiplets identified by each method. Vertical bars (top right) show the sizes of the sets of detected multiplets. If a point is present for a method underneath a bar (bottom right plot panel), this means that this method has identified the particular multiplets that are members of the set denoted by that bar. If multiplet points are present underneath a bar, this means that multiple methods identified the same set of multiplets, i.e. their sets of multiplets intersect. Points are colored by method. Only the 20 largest multiplet sets and set intersections are shown. Note the y axis break.

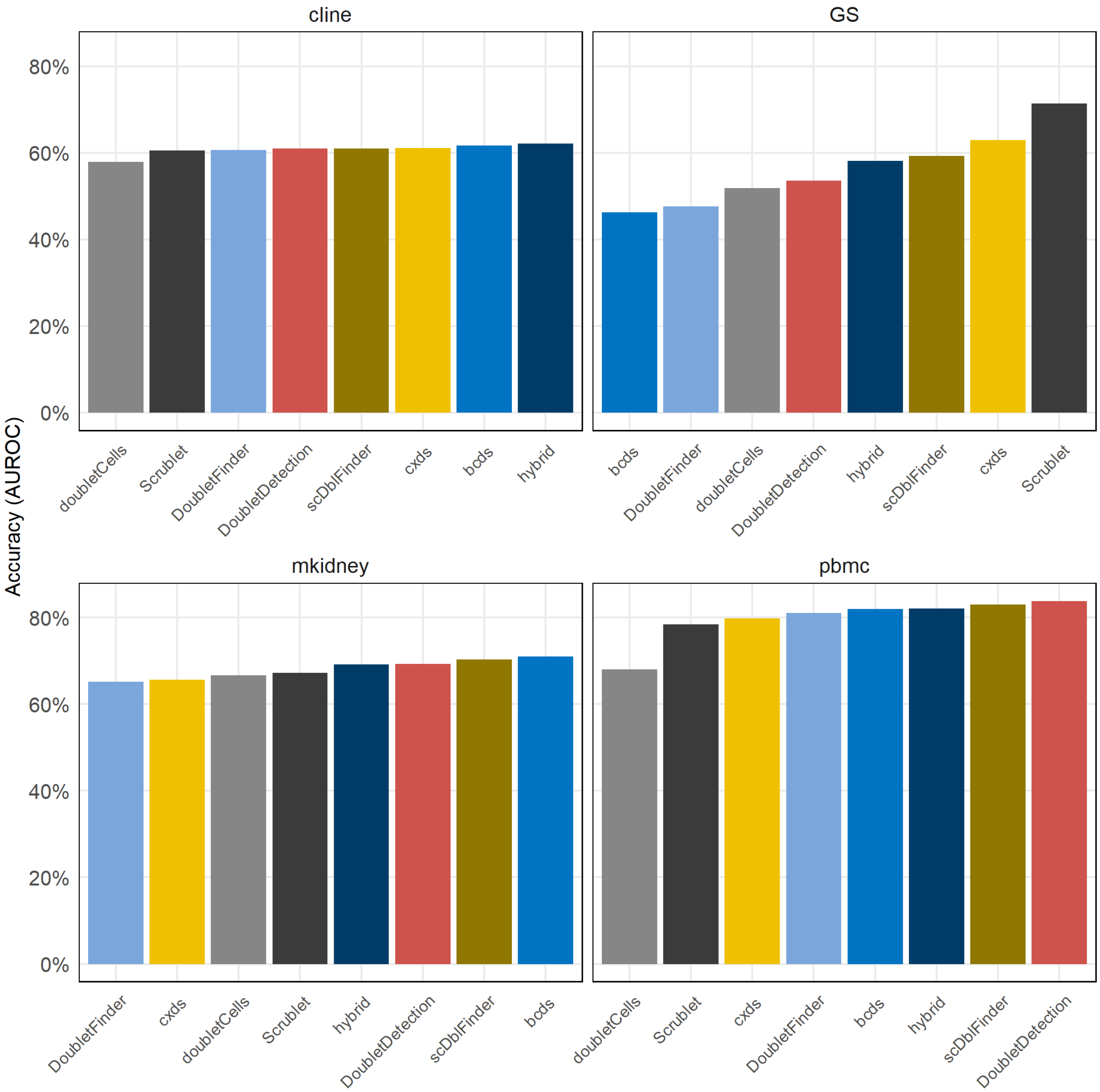


**S4 Fig.**
**AUROC-based accuracy of multiplet detection across datasets and methods**

**S5 Fig.**

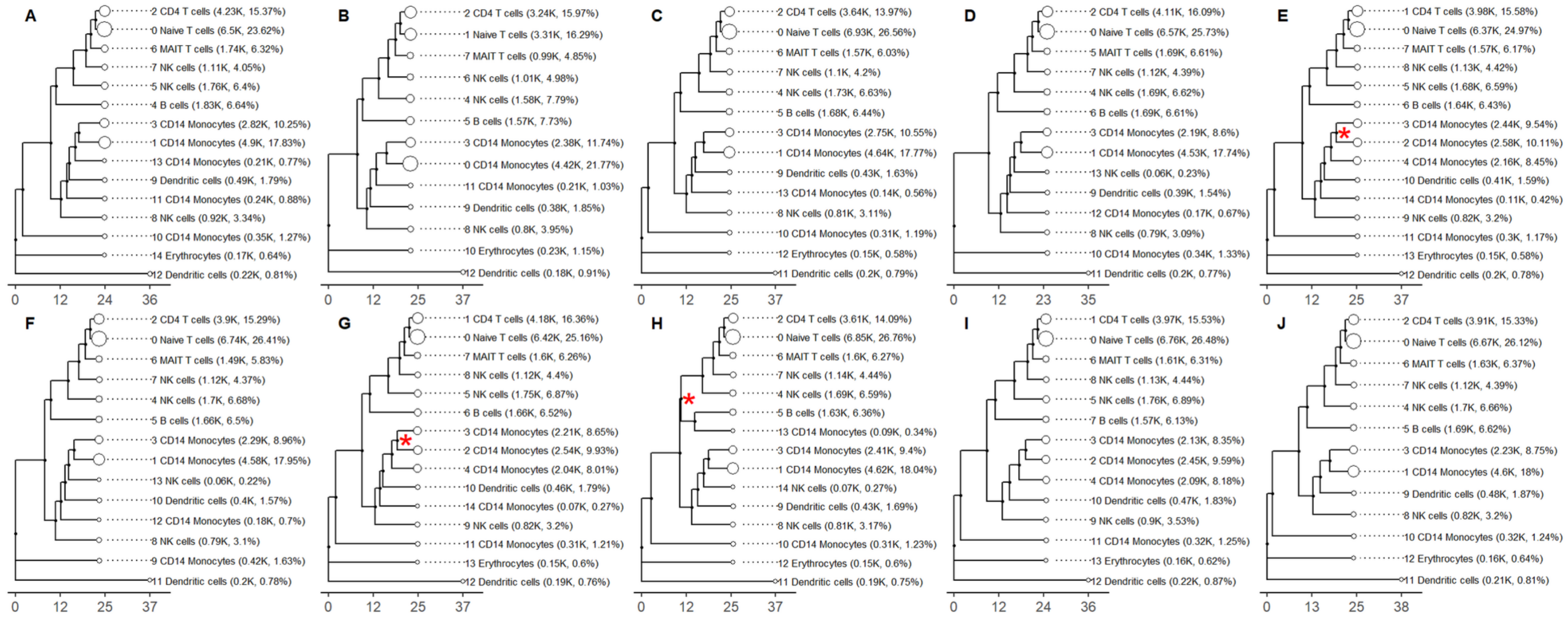


**Bubble tree comparison before and after multiplet removal across all methods**

(A) Gold standard dataset clustered with scBubbletree without removing multiplets. (B) Gold standard dataset clustered with scBubbletree after removing cell-hashing multiplets. (C) Gold standard dataset clustered with scBubbletree after removing DoubletCells multiplets. (D) Gold standard dataset clustered with scBubbletree after removing cxds multiplets. (E) Gold standard dataset clustered with scBubbletree after removing bcds multiplets. (F) Gold standard dataset clustered with scBubbletree after removing hybrid multiplets. (G) Gold standard dataset clustered with scBubbletree after removing scDblFinder multiplets. (H) Gold standard dataset clustered with scBubbletree after removing Scrublet multiplets. (I) Gold standard dataset clustered with scBubbletree after removing DoubletDetection multiplets. (J) Gold standard dataset clustered with scBubbletree after removing DoubletFinder multiplets. The robustness of clustering was assessed with bootstrapping, i.e. by the number of times a cluster emerged out of 1000 bootstrap resamples of the data during clustering. Red asterisks denote the only branches with a bootstrapping value less than 1000.
